# DNA.Land: A digital biobank using a massive crowdsourcing approach

**DOI:** 10.1101/135715

**Authors:** Jie Yuan, Assaf Gordon, Daniel Speyer, Richard Aufrichtig, Dina Zielinski, Joseph Pickrell, Yaniv Erlich

## Abstract

Precision medicine necessitates large scale collections of genomes and phenomes. Despite decreases in the costs of genomic technologies, collecting these types of information at scale is still a daunting task that poses logistical challenges and requires consortium-scale resources. Here, we describe DNA.Land, a digital biobank to collect genome and phenomes with a fraction of the resources of traditional studies at the same scale. Our approach relies on crowd-sourcing data from the rapidly growing number of individuals that have access to their own genomic datasets through Direct-to-Consumer (DTC) companies. To recruit participants, we developed a series of automatic return-of-results features in DNA.Land that increase users’ engagement while stratifying human subject research protection. So far, DNA.Land has collected over 43,000 genomes in 20 months of operation, orders of magnitude higher than previous digital attempts by academic groups. We report lessons learned in running a digital biobank, our technical framework, and our approach regarding ethical, legal, and social implications.

## Introduction

Elucidating the genetic basis of complex traits requires substantial quantities of genomic data ^1^. In the last twenty years, the field has seen an exponential decline in the cost of genomic technologies. As of today, a genotyping array costs on the order of tens of dollars and whole genome sequencing costs thousands of dollars. However, collecting genetic and phenotype data at scale is a time and resource consuming task that poses massive logistical and operational challenges. On top of the costs of genotyping, researchers need to advertise the study, recruit participants, obtain consent, provide DNA collection kits, track and store samples, extract DNA, and prepare the DNA library before data is available in a digital format. Phenotyping requires further resources even if it is done using online questionnaires. These operations are labor intensive and translate to massive costs. For example, the NIH’s Precision Medicine Initiative (“All of US”) has recently allocated $50 million for recruitment centers (“HPO”) and biobank operations that collectively proposed to recruit and handle bio-specimens and basic phenotypic information from a total of 500,000 participants without genotyping. These costs translate to about $100 per participant before the inclusion of more advanced data collection methods such as wearable devices. In addition, the UK Biobank reported that it needed “careful configuration” of its operational chain to support the recruitment of one hundred participants per day in each of its centers ^2,3^.

We sought to develop a cost-effective alternative for collecting genome and phenome data at scale. The past five years have witnessed the advent of large-scale direct-to-consumer (DTC) genetic services for genealogy and personal curiosity, with companies such as 23andMe, MyHeritage, FamilyTreeDNA, and AncestryDNA ^4^. These services provide a dense genotyping array with over half a million SNPs for about $80-$100 per participant. As of today, more than five million individuals have been tested with these services and over ten thousand new DTC kits are purchased daily. With the exception of 23andMe, none of these services collect phenotype information and none of them currently share individual level data with researchers. These policies restrict the ability to migrate data to academic studies by collaborating with these companies. However, all of these services hold the view that the raw genetic information belongs to the tested individual and allow downloading the genomic data in a tabulated textual format. This mechanism provides an opportunity to reach out to individuals to crowd source the raw genetic data for scientific studies, circumventing the cumbersome sample processing procedures of traditional studies.

Previous efforts to crowd source DTC genomic data using an online platform have shown mixed results. For example, OpenSNP.org offers a not-for-profit service and provides a basic mechanism for users to upload their DTC genomic data and publicly share their data, but do not offer features such as privacy controls or a consent to participate in research ^5^. While serving as an important open resource for the community, OpenSNP’s approach has yet to become a viable alternative to traditional genomic data collection. Analysis of uploading dates shows that the website attracts only 1 to 2 participants per day and after five years of operation, it has reached only five thousand participants. Another website for crowdsourcing DTC genomic data is GedMatch.com, which is run by a small for-profit company. This website offers a wide repertoire of genetic-genealogy tools that extend the features offered by DTC companies. By serving the genetic genealogy community, GedMatch has reached critical mass and grown a large community of hundreds of thousands of individuals in approximately five years of operation. However, the website is not focused on basic research: it neither consents users nor collects phenotypic information, and provides no privacy settings, reducing its attractiveness for human genetic research. But its success highlights the possibility of reaching a large scale collection of DTC data by developing a 3^rd^ party service offering added value in the form of genetic-genealogy analysis for participants.

Building upon these observations, we developed DNA.Land, a website to crowd source genomic and phenotypic information for human genetics research. In 20 months of operation, DNA.Land has collected over 43,000 genomic datasets from DTC participants. Importantly, this effort was accomplished by a small team in an academic environment. In this manuscript, we describe the operating guidelines, ELSI approach, and technical details of our website, while highlighting key points and lessons learned to operate a digital biobank. We hope this information can be useful for other academic efforts seeking alternatives to traditional approaches of constructing genetic databases, for start-ups that operate in the growing DTC domain, and for bioinformaticians interested in learning more about the architecture of scalable pipelines for the analysis of genetic data.

## Design principles and user experience

The design and operation of DNA.Land have emphasized two principles: reciprocation and autonomy, which were highlighted by previous studies as a viable route for large scale engagement in genomics ^6–8^. Participants who volunteer their genomic data contribute an essential resource for advancing research. We hypothesized that providing services in return would help maintain user interest and interaction with our study and encourage participation from new users. For every piece of information requested of the user, we aim to reciprocate by displaying online reports detailing interesting information about his or her genome. In addition, we provide a “Learn More” link that explains the value of the information for science and for the user. To respect the autonomy of individuals, we give our users the ability to choose the extent of involvement in the website in terms of data contribution and information sharing.

New users start their interaction with DNA.Land with a short series of steps called the “upload sequence,” which start with account creation and a consent form. Previous studies have shown that users rarely read the terms of service of websites ^9^. To address this challenge, our consent philosophy uses a ‘just-in-time’ presentation of information. Rather than enumerating all possible scenarios as in a traditional consent form for broad research ^10,11^, our consent sets only the framework for the relationships between the user and the study and describes the risks and benefits for sharing genetic data in plain language. While exploring the website, users may decide to increase their involvement by answering questionnaires about health traits or contributing their genealogy data (**Supplementary Note 2**). In these cases, we present additional consent forms that are geared towards the specific feature before allowing the user to contribute more data. The ‘just-in-time’ approach allows the general consent form to be only 1500 words long, or a five-minute read, increasing the chance that users will read it. We share the consent language under CC-BY-2.0 to facilitate adoption by the community (**Supplementary Note 1**). After the consent, participants upload their genetic data and can optionally provide minimal information about themselves. We currently accept data files from 23andMe, AncestryDNA, and FamilyTreeDNA. In the future, we plan to support additional sources and other formats, such as whole exome sequencing.

Once the user has logged in, the main profile page presents three primary types of reports to users: ancestry composition, relative matching, and trait prediction (**Supplementary Figure 1**). To avoid the regulatory complexities involved in sharing health information, traits listed in the trait prediction report are not disease-related and describe only physical and wellness features such as height and neuroticism. Both the trait prediction and relative finder reports implement a ‘just-in-time’ consent for participation, and in the case of the relative finder, users choose to provide a public username and email address visible to other potentially related DNA.Land users; about 90% of our users opted-in for this feature. The trait report, having launched a year after the main site, currently has a 34% participation rate. After completing these steps, users will initially see an “in progress” status on their account homepage as their data is processed. On average, the ancestry reports are available after 7.1 hours (median: 4.6 hours) (**Figure 2A**) and the relative matching and the trait predictions are processed by batch every 12 hours, so typically users will wait a maximum of 24 hours for results.

Interestingly, the popularity of the reports did not match our initial expectations. We initially believed that the ancestry report would generate only secondary interest among users as similar reports are returned by DTC services. However, the ancestry report has proven to be one of the most popular features and generates nearly equal traffic to the relative matching report (**Supplementary Figure 2A**). The launch of a more visually appealing ancestry report in April 2016 generated a massive spike in traffic, and we have since observed many participants publicly sharing their ancestry results in Facebook pages dedicated to genetic genealogy. On the other hand, we believed that users would highly value the option to download their fully imputed genome with 39 million variants compared to their half a million array. We instead found that most users do not have the computational resources to analyze their genome and this feature proved to be infrequently used ^12^.

Finally, we provide tech support and engagement for our users through a dedicated member of our team. The need for this task became apparent when we were flooded with hundreds of emails after the launch, which strained our ability to respond and diverted significant amounts of time from the development team. In addition, our DNA.Land Facebook page has become a place for users to report bugs and pose questions about the website, whereas our initial expectation was that it would only serve for promotional purposes with minor importance. Our tech support answers emails three times a week, responds to user comments on our Facebook page, and writes blog posts promoting DNA.Land on social media, keeping users appraised of our development efforts.

## Data Acquisition during the project

DNA.Land collects many forms of data from users including genome-wide genotyping data, basic demographic information about the participant and his or her immediate family, and questionnaires about traits. With exception of the genomic information, all other types of data are optional for participation in DNA.Land.

We launched DNA.Land on Oct. 2015. As of May, 2017, the project has collected 43,000 genomic datasets from participants. About 45% of the users submit files from AncestryDNA, 40% from 23andMe, and 15% from FamilyTreeDNA. In general, we can divide the participation rates into three phases. The launch phase in the first month saw a rapid rate of growth of nearly 8,000 genomes. Then, after the initial excitement, the rate declined to an average of 900 genomes per month. Finally, after launching the improved ancestry report in April 2016, we have seen a steady growth of nearly 2,000 new genomic datasets per month (**Figure 1**). We also allow users to delete their account at any time. Since the launch of the website, the deletion rate has remained at an average fraction of 4.9% of new user uploads. The reasons for deletions are mostly technical and reflect users that encountered technical problems such as uploading a truncated genome file. In addition, about 6.3% of all submitted the genomes are essentially identical. These cases mostly reflect users that were tested with more than one company and created a separate profile for each one of their genome datasets.

**Figure 1:**
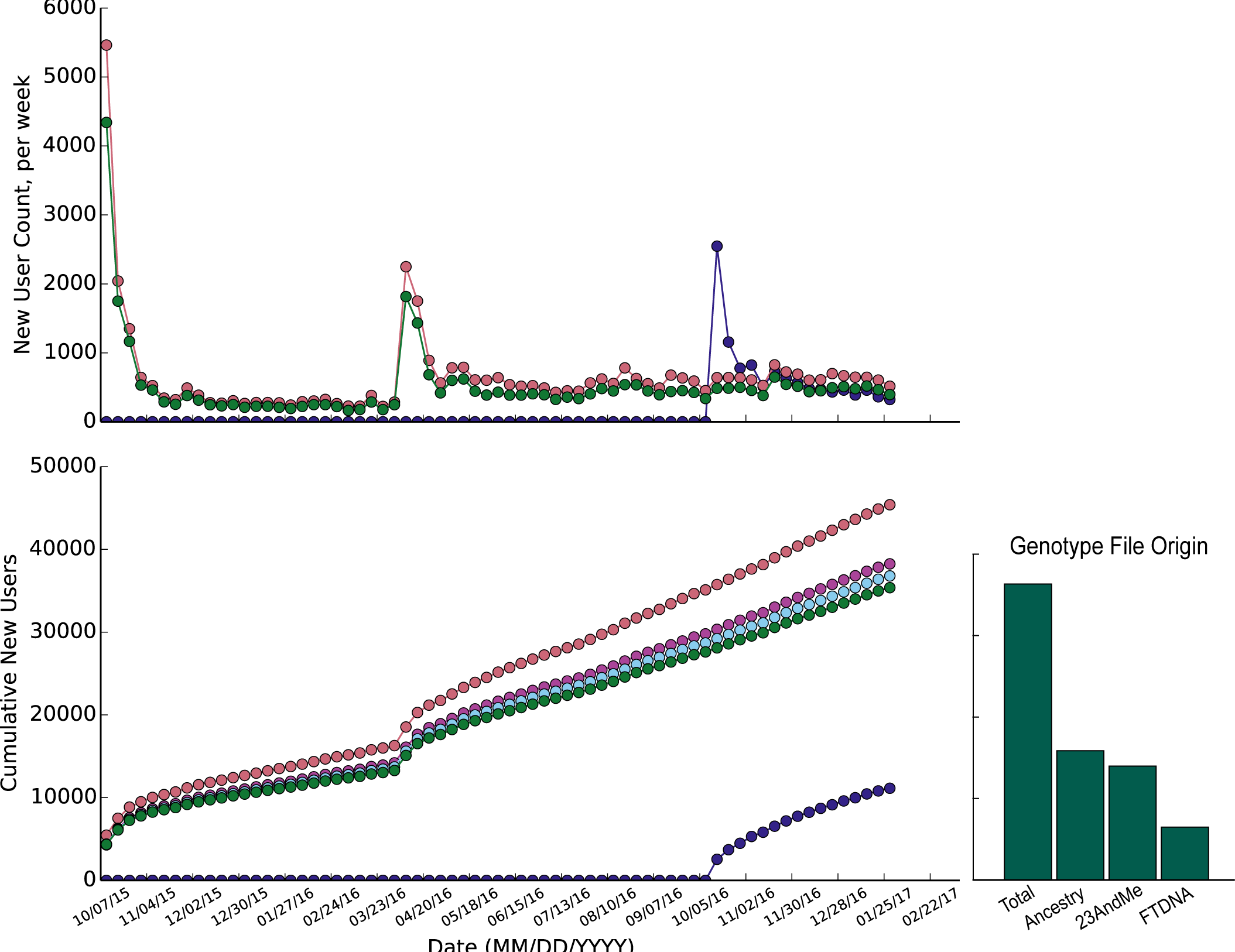
The number of new users participating in DNA.Land since launch on Oct. 9, 2015. Points represent weekly totals, and grid ticks delineate spans of 4 weeks. (Top) New user counts per week. Pink indicates the number of new user registrations. Green indicates the net number of user genomes uploaded, with users who subsequently deleted their accounts subtracted. Dark Blue indicates the number of users consenting to and completing at least one of the trait prediction questionnaires. Large spikes in new user uploads occurred during launch and after the release of an updated ancestry report in April 2016. (Bottom) Cumulative new users per week since launch. In addition to the above, Purple indicates the number of users who submitted email verification of their accounts after registration, and Light Blue indicates the total number of users who uploaded genomes. The bar graph corresponds to the net genomes uploaded (Green) and indicates the proportion of total genomes arriving from each currently-accepted direct-to-consumer genotyping company.

We gather phenotypes by providing users with various questionnaires about physical and health traits (**Supplementary Note 2**). Each questionnaire pertains to a single trait, and users may choose which questionnaires to complete. To facilitate participation, we limited the number of questions in each questionnaire to a maximum of 15, and most users spend less than 2 minutes completing each questionnaire **(Supplementary Figure 3C)**. We launched the questionnaires in October 2016, a year after DNA.Land launched, and since then about 12,000 of our users have completed at least one questionnaire. Users have since answered over 275,000 questions in total, or about 3,100 questions per day since the feature’s launch. We did not discover any significant differences between participation rate in the questionnaires although they sampled very different traits.

We also give users the opportunity to provide detailed information about relatives, with an emphasis on identifying nuclear families. We have integrated into DNA.Land’s relative finder an option for users of Geni.com, a website for building family trees, to link their Geni accounts with those of their matching relatives on DNA.Land. Family trees built by Geni.com users have been shown to facilitate large-scale analyses of populations, such as historical migration patterns ^13^. We also provide survey questions for users to directly identify their mother and father. Lastly, an analysis of the results of the relative matching algorithm across all DNA.Land users shows that 7,100 profiles have at least one immediate family member. Additional information about relative matching statistics of DNA.Land users are presented in **Supplementary Figure 4**.

Analysis of the demographic data provided by users shows that the average participant is of North European ancestry in her late 40s (interquartile region: 36-63 years old) (**Supplementary Figure 4C**). We see a slight over-presentation of self-reported females (53%) versus male (47%). To understand the ethnic composition of our study, we analyzed the genetic ancestry of individuals and identified the leading ancestry component of each individual. While this measure may not directly correspond to how users self-identify their ancestry ^14^, it provides a proxy for the demography of in our data. The genetic analysis shows that the primary ancestry of 53.9% of our users is Northern European, with the next most common groups from other parts of Europe (**Supplementary Figure 5)**.

## Data acquisition costs

DNA.Land employs a hybrid cloud design to reach a cost-effective, scalable operation (**Figure 2C**). **Supplementary Note 3** extensively documents the architecture of the project, and we outline here only general details important to the operation costs. Briefly, the front-end of the website operates on an Amazon Web Services (AWS) EC2 reserved instance. It provides the web interface for managing users, collecting genomic and phenotypic data, compiling surveys, and reporting relative matching and trait prediction results. The pipeline for processing of genomic data (e.g. imputation and ancestry analysis) is executed on AWS spot instances, which process each genome in parallel and allow us to scale out quickly in periods of high demand. The imputation and ancestry results are stored on AWS S3 storage. A physical in-house server then runs relative matching and trait prediction processes, which are CPU,RAM and disk intensive. These processed results, including lists of inferred relatives, are transferred to the database on the front-end server.

**Figure 2:**
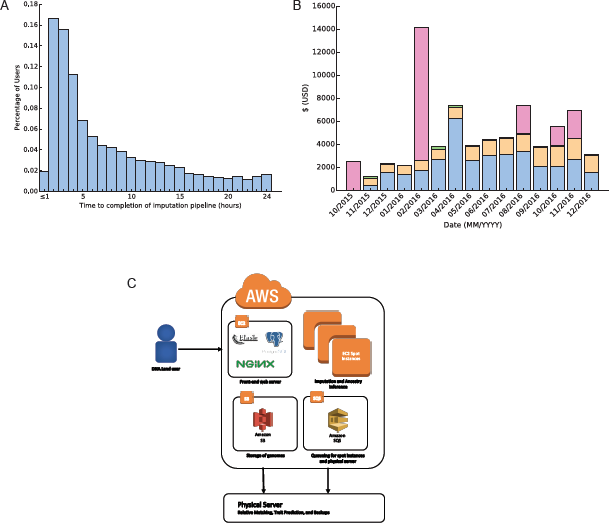
DNA.Land operation and expenses **(A)** The distribution of processing time for newly uploaded genomes in DNA.Land. The processing pipeline involves conversion of the uploaded file to 23andMe format, imputation, and ancestry inference, which occur on Amazon EC2 spot instances. Output files are stored using the Amazon S3 service. **(B)** Monthly expenses for all AWS services. EC2 services (Blue) are used to process new users in the pipeline. S3 (Yellow) is used to store uploaded genotype files and any output files from the imputation pipeline. Transfer costs (Green) pertain to user downloads of their imputed genome files. Irregular costs (Pink) indicate purchases of EC2 reserved instances, as well as purchase of our current physical server, in February 2016. **(C)** Overview of DNA.Land architecture. EC2 spot instances process uploaded genome files and perform imputation and ancestry inference. The physical server performs computation involved in relative matching and trait prediction, and performs backups of genotype data. Genotype files and output files of the impute pipeline are stored in an S3 repository, and selected results are stored in a database on the frontend server. SQS is used to manage assignment of new users to spot instances or processing by the physical server.

The data acquisition costs of our digital approach are low and translate to a few dollars per genome. The costs of running our hybrid cloud operation is on the order of $5000 per month (**Figure 2B**), which includes compute engines, storage, and transfer costs, in addition to irregular costs for development and purchase of our in-house server. To keep the costs low, we have developed an automated bidding system that will bid for spot instances for up to $0.60 per hour, but we can manually decide to bid higher prices in situations of acute need, such as the days following a feature launch during which we experience an influx of new users. As of December 2016, the cumulative cost has been approximately $73.4k, or about $2 per genome-wide genotyping array, a phenome that consist of tens of data points, and genealogy information. In addition, the DNA.Land team has consisted of approximately two full time academic programmers who are mainly required for the development of new features to collect new types of information, and a part-time position for technical support.

## Discussion

We have described a method to gather direct-to-consumer genotype data at low cost and low personnel requirements relative to traditional genotyping methods. In the span of 1.5 years, we have managed to obtain over 43,000 genomes, many of which are paired with additional phenotypic, demographic, and family data.

We credit the success of DNA.Land to several factors. First, we achieved great momentum immediately after the launch of the project, and within the first month of operation we had collected over 8,000 genomes. We attribute the successful launch to working closely with leaders in the genetic genealogy community, who promoted the resource to their social media followers and were paramount to communicating to us the needs of the community. In addition, the initial website already included several interesting features not presented in existing DTC reports, such as a visualization of shared IBD segments between matching relatives, which AncestryDNA does not show. Second, we invested considerable efforts into addressing user concerns on a variety of issues including the quality of our results, privacy and consent policies, and even suggestions for improving our user-interface, such as making our visualizations color-blind friendly. We posit that this process, while resource-intensive, has signaled to the community that we are serious partners that can be trusted with their information. Third, we placed an emphasis on scalable software. After the initial growing pains of stabilizing the website, the day-to-day operation of DNA.Land has required only minimal efforts to maintain. This has allowed our small team to mainly focus on development of new features and reports, which further drive participation. This stands in contrast to traditional studies that involve sample handling and the need to scale personnel to get to the next sample.

We also faced a few challenges in running DNA.Land. First, in academia, the availability of scientific software is usually welcomed regardless of its quality. This is not the case when providing a public website for a non-academic population. Most of our users showed little patience for technical issues in our website, and we found very quickly that we needed to operate on the highest standards of software development and quality assurance, such as support for various browsers as well as mobile and laptop devices. Initially, this led to high stress levels when launching new features and to much longer development cycles than anticipated. We addressed this issue in part by having a development environment that enables prototyping and testing of code before the launch. In addition, we found that soft launching (launching without substantial promotion) to be a more reliable path. This technique has allowed us to test the feature with a smaller set of users and detect technical issues before the feature is discussed and promoted widely on the Facebook groups of our participants. In the future, we hope also to be able to launch a feature only to a small subset of users, but currently our framework does not support this option.

Second, our experience highlights the necessity of a ‘customer support’ function. We were initially overwhelmed by the amount of communication from participants, mainly on our Facebook page which we had established to release messages to the community. We did not anticipate the necessity of a support function before the launch and found ourselves answering thousands of emails and Facebook posts in the first week of operation while managing development issues that occurred during the launch. Answering unhappy participants (some of which are far more critical than “third reviewers”) is not a common task in academic lab and was highly demanding. We encourage others that undertake such a similar endeavor to dedicate a member of the team to answering those emails. In addition, we greatly benefited from an internal system developed by the team that allows tracking the status of each sample in the computational pipeline. This has allowed us to serve participants with accurate information and manage technical issues.

Finally, we learned to pay closer attention to the “actionability” of the data from the user perspective. As researchers, we usually encourage broad and detailed data sharing, but we found some challenges to that philosophy among our participants. For example, we thought that the imputation feature would be of high demand, as we generate for participants the status of 39 million variants from an array which may have only 700,000 SNPs. However, this feature met negative feedback from many users who found that it was not clear what to do with the file and noted that the file is impossible to open with standard applications such as Excel or Notepad. We have addressed this by development of more tools, such as DNA.Land Compass ^12^, which provides a GUI-based website to browse the data and learn about each SNP.

The future for DNA.Land involves more granular consent and expansion of the ways we collect phenotypic information. We are developing a method for participants to share their data with other organizations using an organization-specific consent. In a first attempt, we recently partnered with the National Breast Cancer Coalition (NBCC), a patient advocacy group, to collect genotype and phenotype information for breast cancer research. We re-consent users who participant in our survey and allow them to opt-in for sharing their genome with the NBCC under a specific code of conduct provided by the NBCC. One month after “soft launching” the feature, more than 3,300 users have started the survey, and more than 2,500 (around 80%) have completed it. We aim to create more opportunities that will empower participants to decide for themselves about sharing their data. In addition, we aim to reduce the burden on our participants when collecting phenotypic information. The current procedure of answering questionnaires is cumbersome and does not scale well, as it requires participants to repeatedly visit the website. The last few years have highlighted the rise of digital phenotypes, which refers to quantifying phenotypes from human interactions with digital technology ^15^. Recent studies have shown that a range of traits can be measured with data collected on web activity. These include measuring five big personality traits from Facebook likes ^16^, highly accurate quantification of heart rate from videos ^17^, and finding early signals of pancreatic cancer from Internet searches^18^. Unlike traditional questionnaires, digital phenotypes require less labor from the participant as they leverage existing data using APIs of social media sites such as Facebook and allow measurement of longitudinal changes. We hope to focus on collecting such phenotypes after proper consent from our participants.

As an ultimate goal, we hope to create a digital biobank that integrates streams of data from genetic, genealogical, and social media resources. This approach will establish a complementary effort to existing large-scale traditional studies. Our data-intensive society offers growing numbers of opportunities to harness existing resources, and we envision that the value and scope of such integrative approaches will continue to rise.

## Acknowledgments

Y.E. holds a Career Award at the Scientific Interface from the Burroughs Wellcome Fund. This study was supported by a generous gift from Andria and Paul Heafy to the Erlich Lab, funding from the National Breast Cancer Coalition, and support from Amazon Web Services’ Education Grants. We thank the tens of thousands of DNA.Land participants, especially our early adopters whose feedback was integral in our efforts to improve the site, and genetic genealogist Cece Moore for her valuable advice.

## References

1. Ashley, E. A. Towards precision medicine. Nat. Rev. Genet. 17, 507–522 (2016).

2. Sudlow, C. et al. UK biobank: an open access resource for identifying the causes of a wide range of complex diseases of middle and old age. PLoS Med 12, e1001779 (2015).

3. Downey, P. & Peakman, T. C. Design and implementation of a high-throughput biological sample processing facility using modern manufacturing principles. Int. J. Epidemiol. 37, i46–i50 (2008).

4. Khan, R. & Mittelman, D. Rumors of the death of consumer genomics are greatly exaggerated. Genome Biol. 14, 1 (2013).

5. Greshake, B., Bayer, P. E., Rausch, H. & Reda, J. OpenSNP–a crowdsourced web resource for personal genomics. PLoS One 9, e89204 (2014).

6. Erlich, Y. et al. Redefining genomic privacy: trust and empowerment. PLoS Biol 12, e1001983 (2014).

7. Delaney, S. K. et al. Toward clinical genomics in everyday medicine: perspectives and recommendations. Expert Rev. Mol. Diagn. 16, 521–532 (2016).

8. Wilbanks, J. & Friend, S. H. First, design for data sharing. Nat. Biotechnol. (2016).

9. Bakos, Y., Marotta-Wurgler, F. & Trossen, D. R. Does anyone read the fine print? Consumer attention to standard-form contracts. J. Leg. Stud. 43, 1–35 (2014).

10. Lunshof, J. E., Chadwick, R., Vorhaus, D. B. & Church, G. M. From genetic privacy to open consent. Nat. Rev. Genet. 9, 406–411 (2008).

11. Ball, M. P. et al. A public resource facilitating clinical use of genomes. Proc. Natl. Acad. Sci. 109, 11920–11927 (2012).

12. Curnin, C., Gordon, A. & Erlich, Y. DNA Compass: a secure, client-side site for navigating personal genetic information. bioRxiv 77867 (2016).

13. Kaplanis, J. et al. Quantitative analysis of population-scale family trees using millions of relatives. bioRxiv 106427 (2017).

14. Bryc, K., Durand, E. Y., Macpherson, J. M., Reich, D. & Mountain, J. L. The genetic ancestry of African Americans, Latinos, and European Americans across the United States. Am. J. Hum. Genet. 96, 37–53 (2015).

15. Jain, S. H., Powers, B. W., Hawkins, J. B. & Brownstein, J. S. The digital phenotype. Nat. Biotechnol. 33, 462–463 (2015).

16. Kosinski, M., Stillwell, D. & Graepel, T. Private traits and attributes are predictable from digital records of human behavior. Proc. Natl. Acad. Sci. 110, 5802–5805 (2013).

17. Wu, H.-Y. et al. Eulerian video magnification for revealing subtle changes in the world. (2012).

18. Paparrizos, J., White, R. W. & Horvitz, E. Screening for pancreatic adenocarcinoma using signals from web search logs: Feasibility study and results. J. Oncol. Pract. 12, 737–744 (2016).

